# Predicting Evolutionary Transitions in Tooth Complexity With a Morphogenetic Model

**DOI:** 10.1101/833749

**Authors:** Aidan M. C. Couzens, Karen E. Sears, Martin Rücklin

## Abstract

The extent to which evolutionary transitions are shaped by developmental bias remains poorly understood. Classically, morphological variation is assumed to be abundant and continuous, but if morphogenesis biases how traits vary than evolutionary transitions might follow somewhat predictable steps. Compared to other anatomical structures, teeth have an exceptional fossil record which documents striking evolutionary trajectories toward complexity. Using computer simulations of tooth morphogenesis, we examined how varying developmental parameters influenced transitions from morphologically simple to complex teeth. We find that as tooth complexity increases, development tends to generate progressively more discontinuous variation which could make the fine-tuning of dietary adaptation difficult. Transitions from simple to complex teeth required an early shift from mesiodistal to lateral cusp patterning which is congruent with patterns of dental complexification in early mammals. We infer that the contributions of primary enamel knot cells to secondary enamel knots which are responsible for patterning lateral cusps may have been an important developmental innovation in tribosphenic mammals. Our results provide evidence that development can bias evolutionary transitions and highlights how morphogenetic modelling can play an important role in building more realistic models of morphological character evolution.

## Introduction

Despite more than two centuries of investigation the extent to which ‘laws of variation’(1) govern the tempo and mode of evolution remains poorly understood (2–5). Both as a simplifying assumption and because of ignorance about generating processes, morphological variation has traditionally been assumed to be abundant and continuous (1, 6). However, several decades of evo-devo studies have highlighted how interactions between genes, cells, tissues, and the external environment, favour the generation of some types of variation, whilst conspiring against others (2, 4, 7, 8). This ‘anisotropic’ model of morphological variation implies that evolutionary transitions are more likely to follow some evolutionary pathways than others, and suggests that knowledge of morphogenesis can be used to predict some aspects of morphological evolution (9–11). But, the extent to which these evo-devo insights have penetrated other areas of evolutionary biology, such as phylogenetics, is limited (12–14), at least in part because empirical models of developmental bias are sorely lacking. As an example, the most widely used model of character substitution in morphological phylogenetics, the Markov k (Mk) model (15), assumes equal likelihoods of transition between character states as well as character independence; both assumptions which can be violated by developmental correlations (11, 12, 16, 17). Despite accumulating evidence for wide-spread developmental non-independence and bias in how morphological traits vary (4) there has been limited progress in developing more realistic models for morphological character substitution. At the same time there has been a trend toward ever larger morphological character matrices (e.g. 18, 19) whose underlying properties such as substitution rates and independence remain largely unknown. These concerns have contributed to a widespread scepticism about the reliability of morphological data for building and dating phylogenies (14, 20–22).

Fossil dentitions are often the only phylogenetic link between living and extinct species in evolutionary studies (22–24) and they therefore provide an important morphological system to assess how developmental bias might influence character evolution (9, 11, 25). One of the most pervasive features of dental evolution is the tendency of separate vertebrate groups to evolve complex teeth composed of many cusps (26–28). Perhaps the best documented example is the evolution of complex chewing teeth in mammals and their close relatives (29, 30). In the most basal non-mammalian synapsids the upper and lower cheek teeth are dominated by a single large cusp (31) but in many later mammaliaform groups additional tooth cusps were added predominantly along the longitudinal axis, and eventually in a triangular pattern (29). In ancestral therian mammals, increasing the acuteness of this triangular cusp configuration, was a key step in patterning a more rectangular lower molar with a larger crushing basin on the rear of the tooth (32). Once established, iteration of this core lateral cusp pattern offered a pathway for later mammalian groups to increase molar tooth complexity (33). In stem therian mammals, the shift from a longitudinal to almost rectangular lower molar marks the advent of ‘tribospheny’, where an upper molar cusp, the protocone, crushes food within the enlarged talonid basin on the rear of the opposing lower molar (30). Tribopsheny may have evolved separately in the ancestors of monotremes and therian mammals (34, 35), in each case probably involving a similar reorganisation of longitudinally arranged cusp into an increasingly acute triangular configuration.

Despite the diversity of mammalian tooth shapes, most of the differences between mammalian species reflect relatively small changes in the number, spacing and size of tooth cusps (36). The position and size of tooth cusps is predicted by the expression of embryonic gene signalling centres called enamel knots (33, 36) which secrete signalling molecules that regulate growth and folding of the dental epithelium, which ultimately creates tooth shape. Molecular signals from the enamel knot locally inhibit differentiation of new knot cells which together with global growth of the dental epithelium, helps determine the spacing and number of tooth cusps (36). Because the inhibitory zone established by the primary enamel knot strongly influences the size of secondary enamel knots, the number of tooth cusps and thus overall tooth complexity is likely to be closely linked to developmental events that regulate early cusp patterning (37). Computer modelling of these interactions recovers enamel knot patterns similar to basic mammalian tooth types (37–39) but it remains unclear whether such morphogenetic models can predict transitions between tooth types.

Here, we use a computational model of tooth morphogenesis, ToothMaker (38), to investigate: **(1)** how does morphogenesis influence variation in tooth complexity?; and **(2)** does morphogenesis bias evolutionary transitions in tooth complexity? Our results provide insights into the developmental assembly of the mammalian dentition and highlight the potential of morphogenetic models of organ development for building more realistic phylogenetic models of character evolution.

## Results

### Structure of Tooth Complexity Parameter Space

Simulations of tooth development under six different parameter combinations (Fig. 1A) produced large variations in tooth complexity (Fig. 2). Simulations reveal a progressive increase in tooth complexity (measured as occlusal patch count) through developmental time, quantified as the number model iterations (Fig. S1). This increase in tooth complexity was primarily driven by the longitudinal addition of EKs as the epithelium elongates. Developmental trajectories in tooth complexity were generally similar between tooth types except for teeth with laterally patterned EKs which diverge early from other tooth types. Tooth complexity quantified as the number of EKs varied between one and 28, and increases in tooth complexity were generally ordinated diagonally across parameter space. The largest increases in enamel knot number were produced by varying parameters regulating molecular activators and inhibitors or their diffusion rate rather than epithelial growth (Fig. 2). Increasing tooth complexity is coupled with a general increase in the median morphological distance between adjacent teeth in the parameter spaces (Fig. S2). This relationship holds for most parameter combinations irrespective of whether morphological distance is computed using enamel knot number or orientation patch count.

**Fig. 1.**
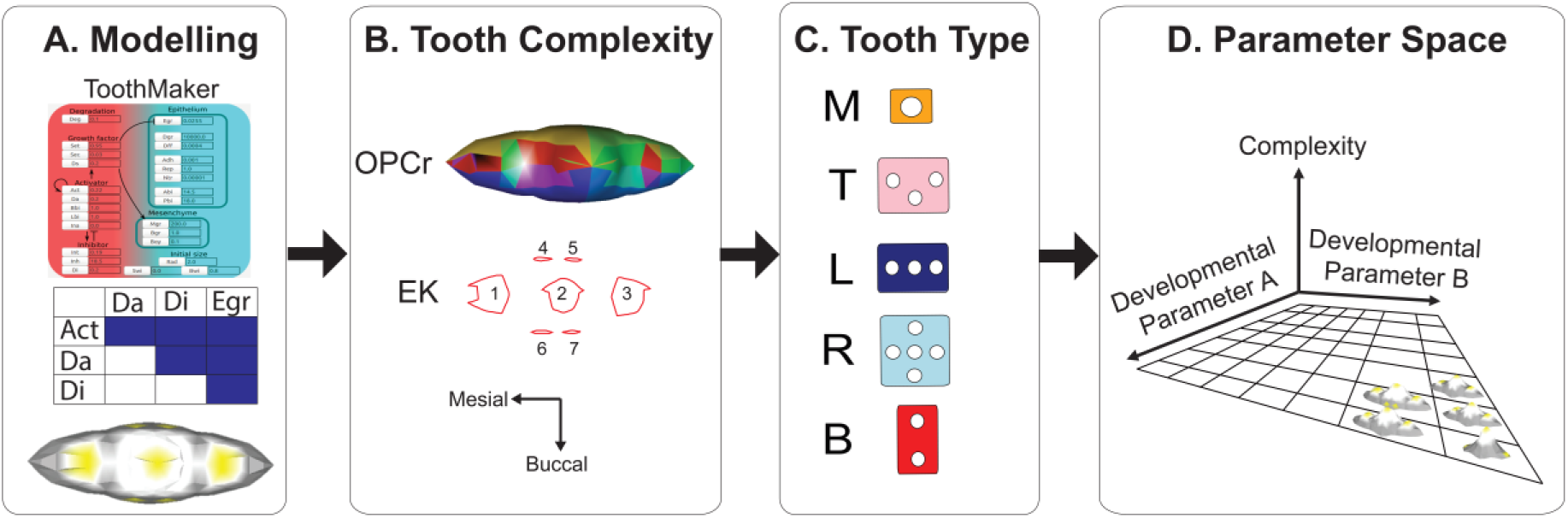
Workflow for construction of parameter space. (*A*). Parameters are varied within ToothMaker for each possible pairwise combination of cellular and signalling factors. Parameter abbreviations: Act, activator autoactivation; Da, activator diffusion rate; Di, inhibitor diffusion rate; Egr, epithelial growth rate. (*B*) Tooth surface complexity is measured using orientation patch count rotated (OPCr) and by quantifying the number of differentiation zones representing enamel knots (EKs). (*C*) Tooth-type was categorised based on occlusal arrangement of EKs into five groups: B, EKs are laterally (buccolingually) separated; L, EKs added longitudinally; ‘T’, EKs form a triangle; R, EK’s added radially, and M, single EK only. (*D*) The parameter space is populated with ‘complexity’ and ‘tooth type’ measurements to generate a developmental morphospace.

### Tooth Type, Complexity, and Transition Rates

Five different tooth types were recognised from simulations based on the organisation of EKs in occlusal view (Fig. 1C). Teeth with the lowest complexity were monocusped teeth (‘M’) with only a single EK, whereas all complex teeth (> 10 EK) had laterally separated knots (Fig. 2). The most commonly generated tooth type based on grid cell frequency were teeth with laterally separated knots (‘B’) and the least common were teeth with radially arranged knots (‘R’). Tooth type and complexity were closely associated across parameter space, with the most complex teeth having either triangular or laterally arranged EKs (Fig. 2). Teeth with laterally separated EKs had tooth complexities that were about double that of other tooth types (Fig. 3).

**Fig. 2.**
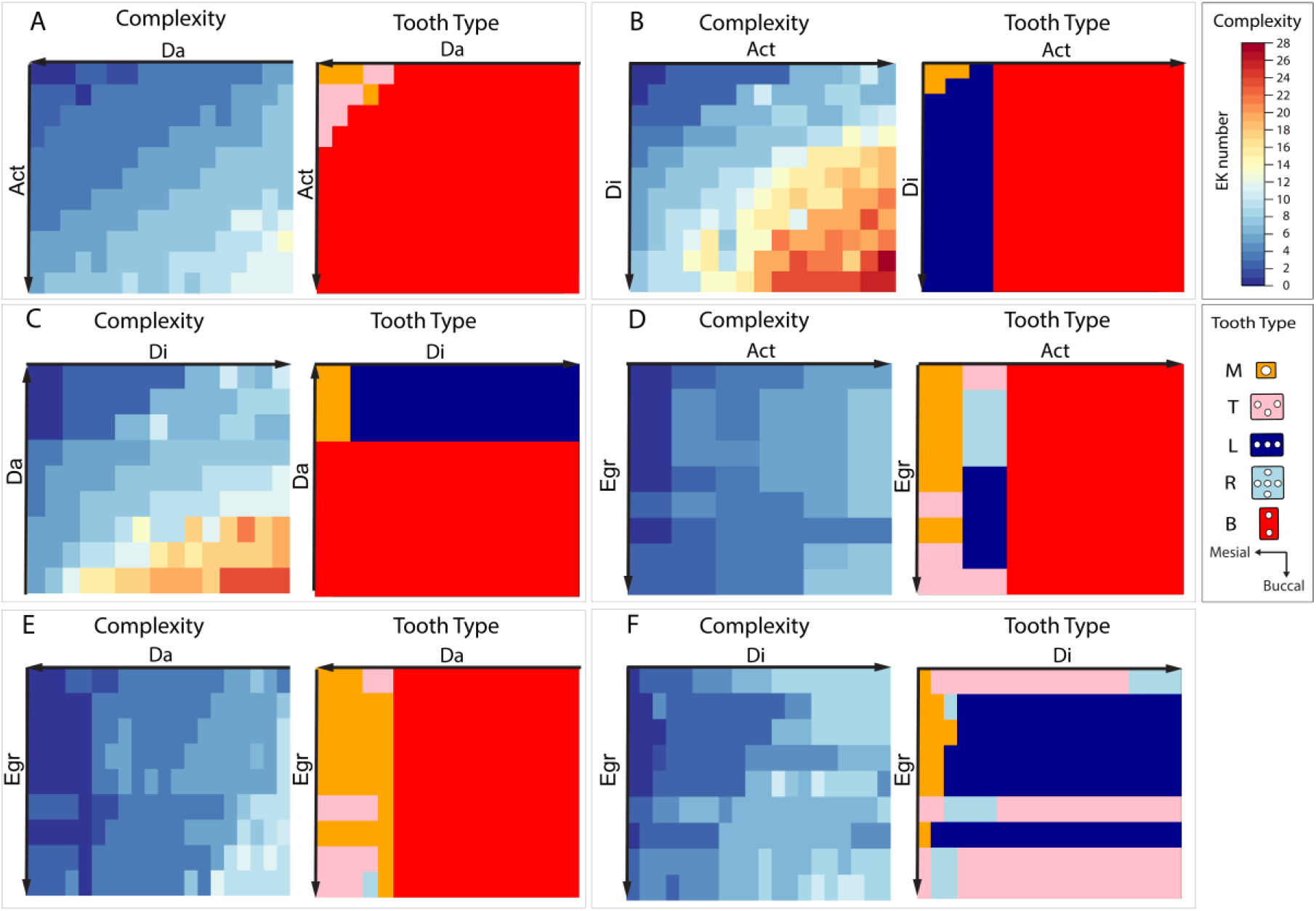
Heatmap representation of tooth complexity and tooth type across developmental parameter space. Parameter space is ordinated with simplest to most complex teeth from top left to bottom right. Axis arrows denote direction of increasing parameter values. Tooth complexity is measured as enamel knot (EK) number. Parameter combinations are as follows: (*A*) Activator autoactivation (Act) and activator diffusion rate (Da). (*B*) Activator autoactivation (Act) and inhibitor diffusion rate (Di). (*C*) Activator diffusion rate (Da) and inhibitor diffusion rate (Di). (*D*) Epithelial growth rate (Egr) and activator autoactivation (Act). (*E)* Epithelial growth rate (Egr) and activator diffusion rate (Da). (*F)* Epithelial growth rate (Egr) and inhibitor diffusion rate (Di).

**Fig. 3.**
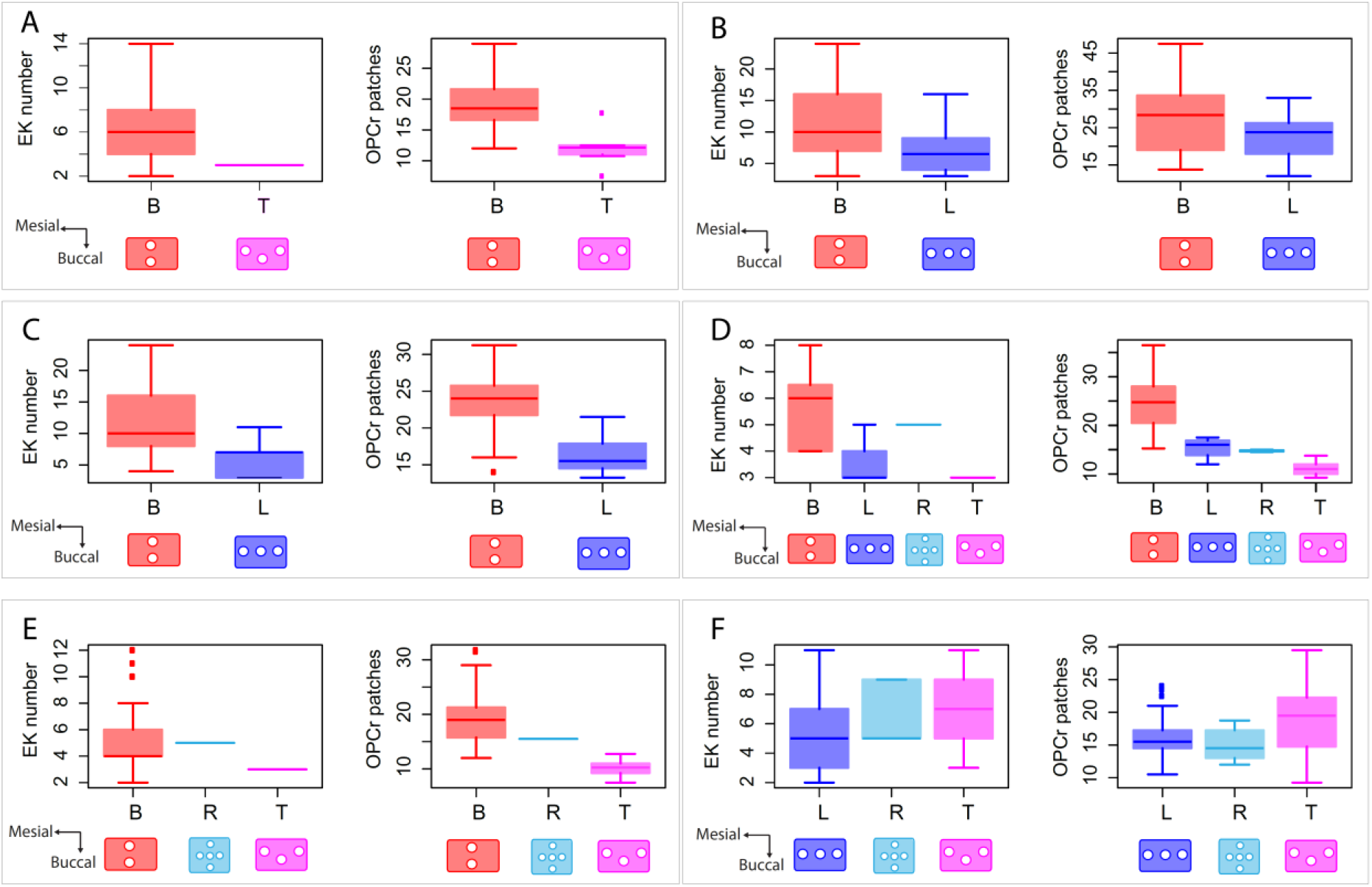
Relationship between tooth complexity and tooth type. Tooth complexity is measured as enamel knot (EK) number and orientation patch count rotated (OPCr). (*A*) Activator autoactivation (Act) and activator diffusion rate (Da). (*B*) Activator autoactivation (Act) and inhibitor diffusion rate (Di). (*C*) Activator diffusion rate (Da) and inhibitor diffusion rate (Di). (*D*) Epithelial growth rate (Egr) and activator autoactivation (Act). (*E*) Epithelial growth rate (Egr) and activator diffusion rate (Da). (*F*) Epithelial growth rate (Egr) and inhibitor diffusion rate (Di).

Transition rates estimated using a ‘shortest-unique-path’ from a simple to complex tooth were uneven, with most transitions from an ‘M’ to ‘B’ tooth occurring by a ‘T’ intermediate (Fig. 2). The proportion of shared perimeter between tooth types was also uneven (Table 1) with ‘L’/ ‘B’ and ‘M’/ ‘T’ tooth types especially likely to share borders. Teeth with an ‘M’ type EK arrangement were twice as likely to share a border with a ‘T’ tooth compared with an ‘L’ or ‘B’ pattern. The ‘R’ type pattern had the most even proportion of shared borders, but this number was low owing to the rarity of this tooth type. Teeth with EKs added mesiodistally (an ‘L’ or ‘T’ tooth) had a three-fold higher probability of bordering a ‘B’ tooth compared with other tooth types.

**Table. 1.**
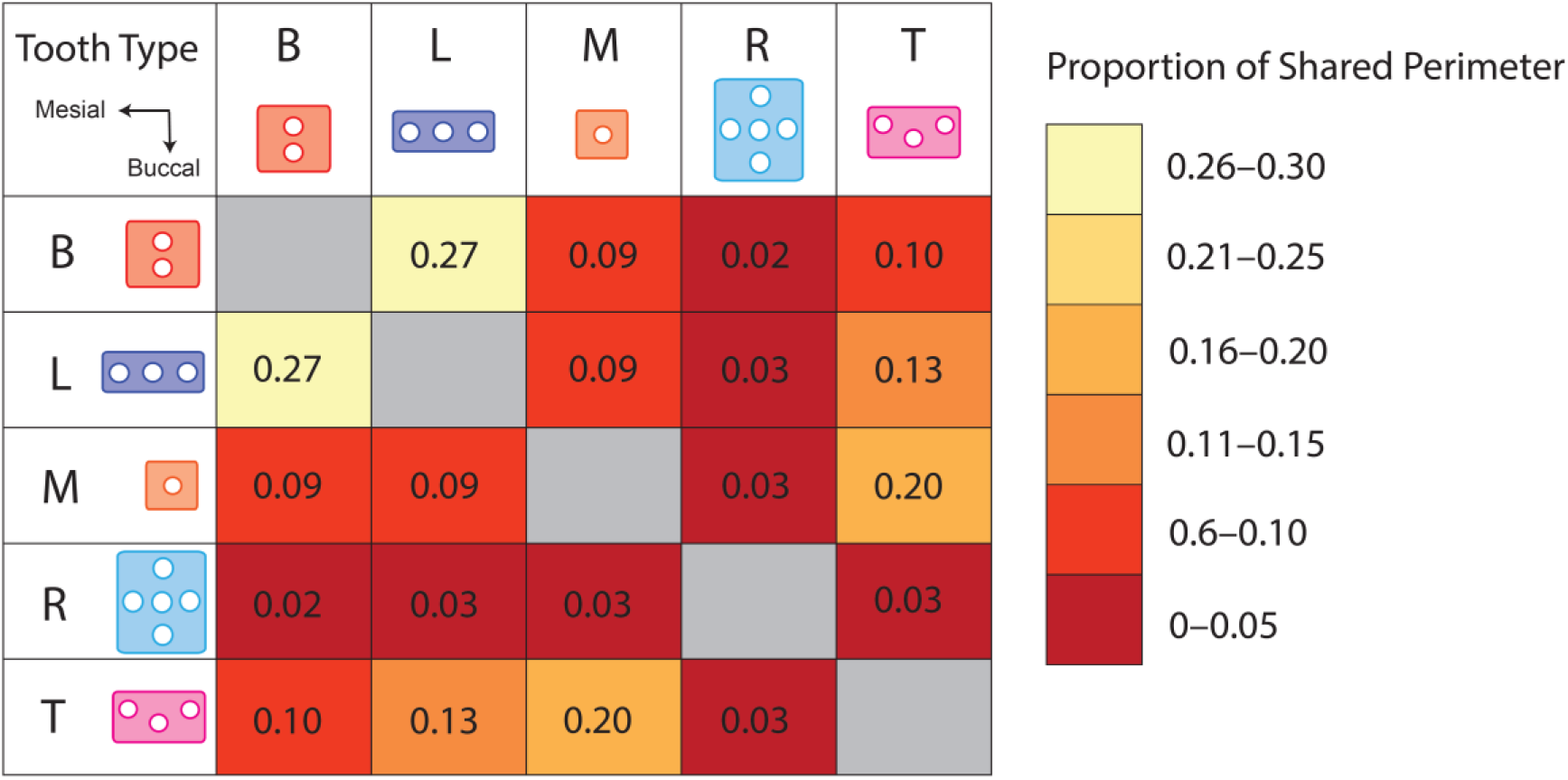
Proportion of shared perimeters between tooth types. Proportions are averages across all parameter combinations.

## Discussion

Developmental processes have long been implicated as a potential source of ‘bias’ or ‘constraint’ in morphological variation but we largely lack explicit models of how this bias might affect evolutionary change (13, 25). Here, we have used a computer model of tooth morphogenesis, ToothMaker, to examine how development biases tooth complexity evolution. Despite varying different types of developmental parameters, we find that complex teeth emerge in all six cases (Fig. 2). Additionally, the way variation in tooth complexity is structured within developmental parameter spaces is similar, with simple and complex teeth being partitioned into distinct regions. In general the increase in tooth complexity is ordinated along a diagonal gradient which supports experimental evidence that gains in tooth complexity are achieved primarily by synchronously changing multiple developmental variables (26). Tooth complexity was not evenly distributed across parameter space and instead zones with more complex teeth had more discontinuous patterns of variation than did those with simpler teeth (Fig. S1). The more ‘rugged’ developmental landscape of complex teeth suggests that complex dentitions are more likely to be developmentally biased than simpler teeth and potentially more difficult to adaptively ‘fine-tune’ (40). However, several highly successful mammalian groups including the multituberculates (41) and rodents (42) evolved very complex teeth (>10 cusps) suggesting that such constraints are not prohibitive, perhaps because these groups acquired gene regulatory innovations that promoted more continuous patterns of dental variation. Most other mammal groups possess simpler teeth, at least in terms of tooth cusp number (43), and their dentitions may have been correspondingly more straightforward to adapt. Additionally, while enamel knot patterning offers a straightforward way to increase complexity, some primate and kangaroo lineages found other ways to increase dental complexity, such as by crenulating the enamel (25, 44), and in ungulates by forming recesses in the enamel called infundibula (45). Changes in non-patterning aspects of the genetic cascade, like mineralization genes, might therefore provide an important route to constraints arising from the tooth cusp patterning cascade.

Tooth complexity was closely linked to specific tooth cusp architectures (Fig. 3) suggesting that, although tooth complexity can be uncoupled from cusp patterns (42), some types of cusp arrangement are biased toward generating more complex teeth. For instance, teeth with laterally arranged cusps similar to the metaconid and protoconid arrangement of tribosphenic mammals had complexities approximately double that of other tooth types (Fig. 3). Teeth with longitudinally arranged cusps typically had no more than six cusps which parallels patterns of tooth cusp number variation along the jaw in pinnipeds and early mammals like *Morganucodon* and *Kuehneotherium* (36, 46, 47). Differences in tooth complexity between teeth with different cusp architectures might be linked to the way that tooth bud growth influences the initiation of secondary enamel knots. For instance, the initial laterally-directed patterning associated with tooth shapes that grow in both lateral and longitudinal axes, creates space obliquely for secondary knots to initiate. In contrast, when growth is longitudinally-directed, new knots can only initiate longitudinally. Comparison of enamel knot patterning ‘histories’ between different tooth types shows that these early patterning events, which determine the basic cusp architecture, drive tooth complexity along very different trajectories (Fig. S2). Thus, whereas variation in the tooth cusp number has been linked to changes in the size of the primary enamel knot in the predominantly longitudinal teeth of seals (36) our findings suggest that in teeth with a stronger buccolingual growth component, tooth complexity becomes more closely correlated to the initial spatial patterning of the secondary knots.

Multiple independent increases in cheek tooth complexity occurred in early mammal evolution as exemplified by the dental evolution of multituberculate (41), australosphenidan (34), and early therian mammals (29). In the ancestors of therian mammals a rectangular lower molar evolved, consisting of a posterior crushing basin, the talonid, which occludes with the upper molar protocone cusp (29, 30). This ‘mortar and pestle’ configuration, a functional hallmark of tribospheny, permitted food to be crushed and sheared in the same chewing stroke (29, 32). During the evolution of therian tribospheny, the ancestrally longitudinal arrangement of tooth cusps on the lower molars of pretribosphenic mammals was modified into an increasingly acute triangular configuration that created a more transversely orientated trigonid wall (Fig. 4B). In early therians like *Ambolestes* and *Kielantherium* the protoconid was displaced buccally and the metaconid somewhat lingually. This brings these cusps almost directly lateral from each other and created a more rectangular lower molar with space for a larger talonid basin (Fig. 4B). Using ToothMaker, we find that in longitudinally patterned teeth similar to pretribosphenic mammals, the secondary knots form from distinct region of dental epithelium, whereas laterally patterned cusps originate within the spatial domain of the primary knot. Lineage tracing of primary knot cells in mouse indicate that primary knot cells contribute to the buccal (e.g. protoconid) (48) and possibly the lingual secondary knot (49). However, it remains uncertain whether the differential contribution of primary knot cells to these secondary centres may influence the latter’s signalling dynamics (50). Nevertheless, the modelling results highlight how the contribution of the primary knot cell population to laterally-positioned secondary knots (Fig. 4B) may have been a key developmental innovation in the origins of tribospheny. Similar laterally separated cusp arrangements evolved independently amongst monotremes, multituberculates, and even amongst Mesozoic crocodyliforms (e.g., *Chimaerasuchus paradoxus*) (27) suggesting that this switch in the cusp patterning axis may have been relatively evolvable (Fig. 4B). Still, the exact developmental processes which prompt the initial shift from longitudinal to lateral cusp patterning remain unclear. Recent evidence suggests that relaxing lateral compressive forces during cusp patterning can alter tooth cusp patterns on rodent molars (51). This raises the interesting question of whether increased developmental separation between the jaw bone and molar teeth during cusp patterning might have been crucial to the origins of therian tribospheny.

**Fig. 4.**
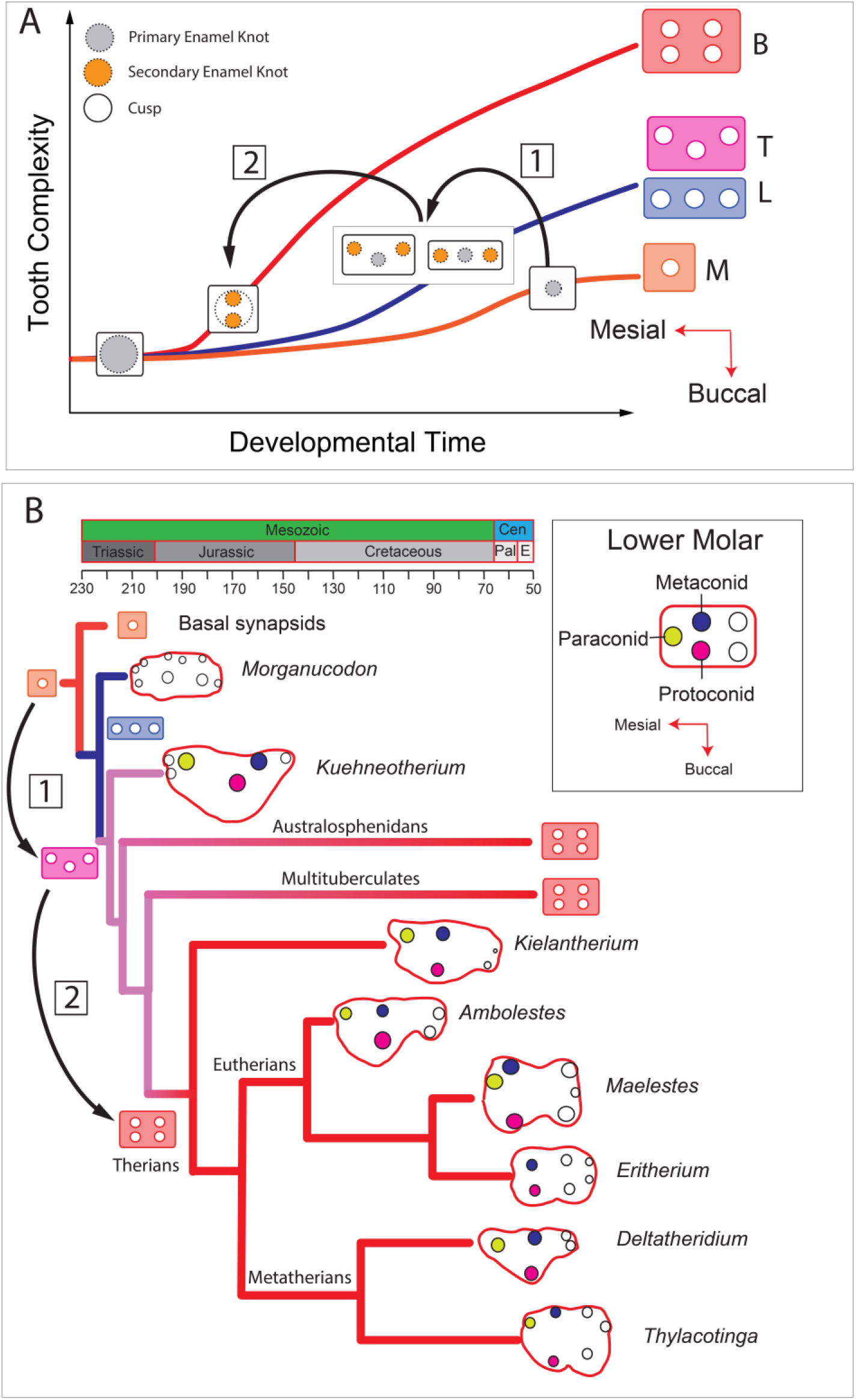
(*A*). Model of cusp patterning transition associated with the origin of complex teeth. (*B*). Schematic evolutionary model of lower molar (M_2_) cusp pattern evolution in early mammaliaforms. Inferred tooth types are illustrated at key nodes. Phylogeny modified after (35).

Mapping the distribution of tooth types across the parameter spaces shows that transitions between different tooth type ‘states’ is uneven. For instance, the transitions with the fewest steps between a simple and a complex tooth involved first adding cusps either longitudinally or in a triangular pattern (Fig. 2). A similar pattern is found if the proportion of grid cell borders shared between tooth types is computed (Table 1). Monocusped and laterally patterned teeth have >70% chance of bordering teeth with a triangular or longitudinal cusp pattern. These findings suggest that complex teeth may be developmentally biased toward evolving by **(1)** adding cusps mesiodistally and then **(2)** laterally (Fig. 4A). This model of character state transition broadly fits with the leading phylogenetic hypotheses of early mammal evolution (35, 52, 53) (Fig. 4B). However, most models of early mammal dental evolution recover a very specific shift from a longitudinal to triangular cusp pattern on the upper and lower molars (35, 52, 53) (Fig. 4B), whereas the developmentally-inferred transition rates are more ambiguous. For instance, the developmental model suggests that a single cusped tooth can readily transform to a triangular state without a longitudinal intermediate. Likewise, transformations from a longitudinal to a buccolingual state were up to three-times as likely as from a triangular state. These incongruencies may reflect the fact that some transitions between tooth types are less developmentally biased than others, perhaps because functional factors are more influential. Additionally, relatively small developmental differences such as lateral growth biases (51) may be all that is required to shift between longitudinal and triangular tooth types.

The reliability of morphological phylogenies has been criticised on a range of fronts but especially due to character non-independence and uncertainties about the correct model of character evolution (11, 20–22). The transition rates estimated between tooth types based on the simulation of some developmental parameters here do not support equal transition probabilities between states as assumed under the Mk model (15). But, a much more exhaustive sampling of the ToothMaker parameter space is needed, to precisely estimate transition rate probabilities as well as their symmetry. Additionally, while we have focused on perhaps the most basic aspect of tooth shape, the cusp pattern, there is a need to consider more phylogenetically relevant dental characters. One possibility is that transition rate parameters estimated from morphogenetic models like ToothMaker could be implemented as a transition rate prior on a traditional Mk model (54) or by using a threshold model where transitions between characters depend on a continuous liability distribution inferred from the parameter space (55). Ultimately, developmental bias might actually prove an asset to morphological phylogenetic inference because trees containing developmentally ‘forbidden’ character state transitions could be excluded out-of-hand. Morphogenetic models also provide a new method to test assumptions about developmental character independence which can help to avoid the problems of high type one error rates in character correlation tests (56). In summary, improved integration between morphogenetic modelling of different organ systems and morphological phylogenetics holds promise for improving our ability to reconstruct the history of life.

## Materials and Methods

### ToothMaker Simulations

Using ToothMaker v.0.6.3 (http://dead.cthulhu.fi/ToothMaker) tooth shapes were simulated by varying four developmental parameters (Tab. S1). Due to the large range of possible parameters (26), four parameters with well-defined developmental roles were selected. These parameters have previously been closely linked to natural variation in pinniped tooth shape (38) and implicated in the patterning of different mammalian tooth types (37). These developmental parameters were: epithelial growth rate (Egr), activator autoactivation (Act), activator diffusion rate (Da) and inhibitor diffusion rate (Di). Pairwise combinations of each were used to construct six unique parameter spaces. The ToothMaker program was implemented with a virtual Linux machine (Ubuntu 16.04.3) mounted on a 64-bit Windows host using the software VMware Workstation 15 Player. The dimensional variables for each developmental parameter were varied from an ‘ancestral’ tooth composed of single cusp. The parameter range over which each developmental factor was varied was determined based on capacity to generate a tooth containing at least 10 cusps (Tab. S1). Inhibitor diffusion rate was allowed to vary over a wider range than other developmental parameters given experimental evidence that diffusion rates for proposed tooth inhibitors like fibroblast growth factor protein can vary more than 13-fold (57). Step sizes for each parameter were varied 2–7 % of the maximum parameter value (Tab. 1). Using the ‘scan parameter’ function the tooth surface from each time point spanning the initial stage (0 steps) to final stage (14000 steps) was exported at 500 step intervals (28 time points in total) in *.off file format.

### Mesh Processing

For each model time point, the *.off grid file containing the cell surface geometry was converted to *.ply surface file in `R` (58) using the `vcgPlyWrite` function in the package `rvcg` (59). Non-manifold vertices were removed and vertex normals coherently aligned using the `vcgClean` function and the surface reflected about the z-axis to invert the vertex normal using the `mirror` function.

### Tooth Complexity and Tooth Type

Tooth complexity was quantified by measuring surface complexity using two methods, orientation patch count rotated, and enamel knot (EK) number (Fig. 1*B*). Orientation patch count (OPC) (42) groups contiguous regions of a surfaces with orientations falling within eight directional bins into patches and then sums the number of patches over the surface. Surfaces with more differently orientated regions will have a higher patch count. A modification of OPC, called OPCr attempts to accommodate for differences in specimen orientation by rotating the surface between 0 and 45°. The OPCr analysis was implemented using `molar_Batch` in the `R` package molaR(60) with `OPCr_steps =` 4, `OPCr_stepSize=` 5.625, `OPCr_minimum_faces =` 3 and all other parameters run as default. Enamel knot number was computed using a macro in Image J (1.52a) that thresholds regions of cell differentiation from occlusal snapshots exported from ToothMaker and quantifies the number of thresholded ‘islands’. All tooth models were categorised into five basic groups based on EK arrangement in occlusal view which we refer to as ‘tooth type’ (Fig. 1*C*). Tooth type categories were: ‘B’, EKs are buccolingually separated; ‘L’, EKs arranged longitudinally; ‘T’, EKs in a triangle; ‘R’, EKs added radially; and ‘M’, single EK only.

### Morphospace

Tooth complexity measurements (based on OPCr and EK scores) and tooth type classifications were mapped onto the parameter space of each parameter combination (Fig. 1*D*). To measure the ‘ruggedness’ of developmental morphospace, a proxy for developmental constraint, the difference in tooth complexity between neighbouring teeth within the parameter space was quantified using a ‘sliding-window’ algorithm written in R. This algorithm computes the median difference in complexity scores between each grid cell in the parameter space (a simulated tooth) and all its adjacent neighbours. To avoid an edge effect, grid cells bordering the margins of the parameter spaces were excluded from the calculation. The equation for the ruggedness calculation is:

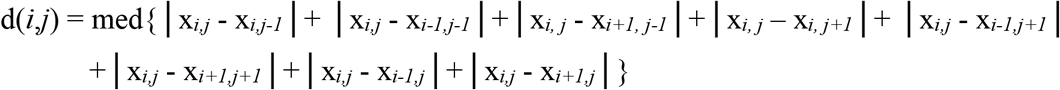

Where:

d is morphological difference in complexity between a grid cell positioned at column *i* and row *j* and all its neighbours.
x is the complexity score at column *i* and row *j*.

## Acknowledgements

We thank J. Jernvall, I. Salazar-Ciudad, T. Häkkinen, and O. Stenberg (Univ. Helsinki) for assistance implementing ToothMaker. P. Gill (Univ. Bristol) and A. Evans (Monash Univ.) provided helpful discussions. R. Beck (Univ. Salford) and B. King (Naturalis) are thanked for feedback which greatly improved the manuscript. This project was supported by NWO, Vidi 864.14.009 grant to M. Rücklin.

---

**Fig. S1.**
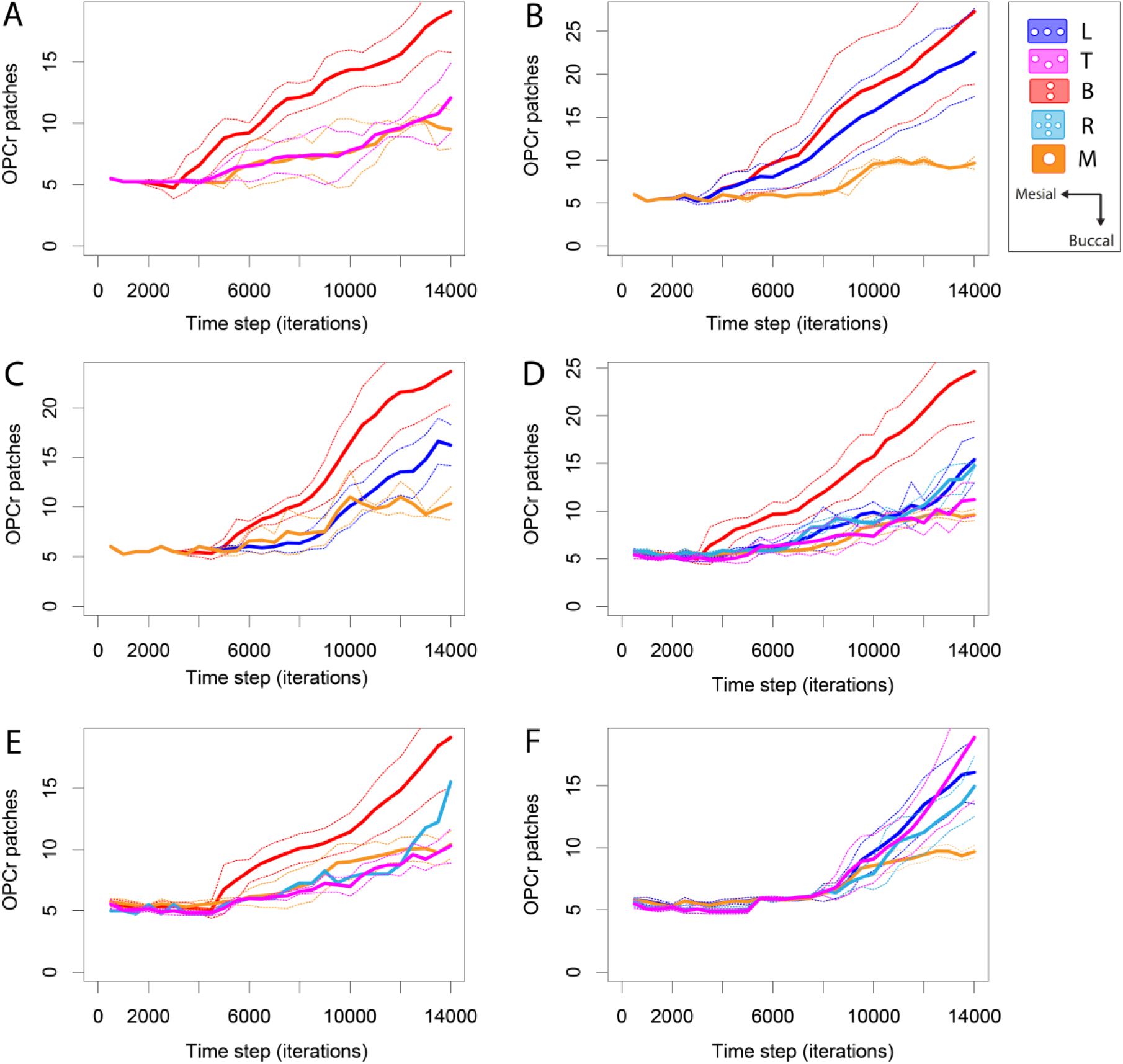
Developmental changes in tooth complexity for tooth types. Tooth complexity is measured as OPCr scores. Confidence intervals represent by dashed lines represent one standard deviation. Time steps represent the number of model iterations ranging from the start (0) to the end (14000). (*A*) Activator autoactivation (Act) and activator diffusion rate (Da). (*B*) Activator autoactivation (Act) and inhibitor diffusion rate (Di). (*C*) Activator diffusion rate (Da) and inhibitor diffusion rate (Di). (*D*) Epithelial growth rate (Egr) and activator autoactivation (Act). (*E*) Epithelial growth rate (Egr) and activator diffusion rate (Da). (*F*) Epithelial growth rate (Egr) and inhibitor diffusion rate (Di).

**Fig. S2.**
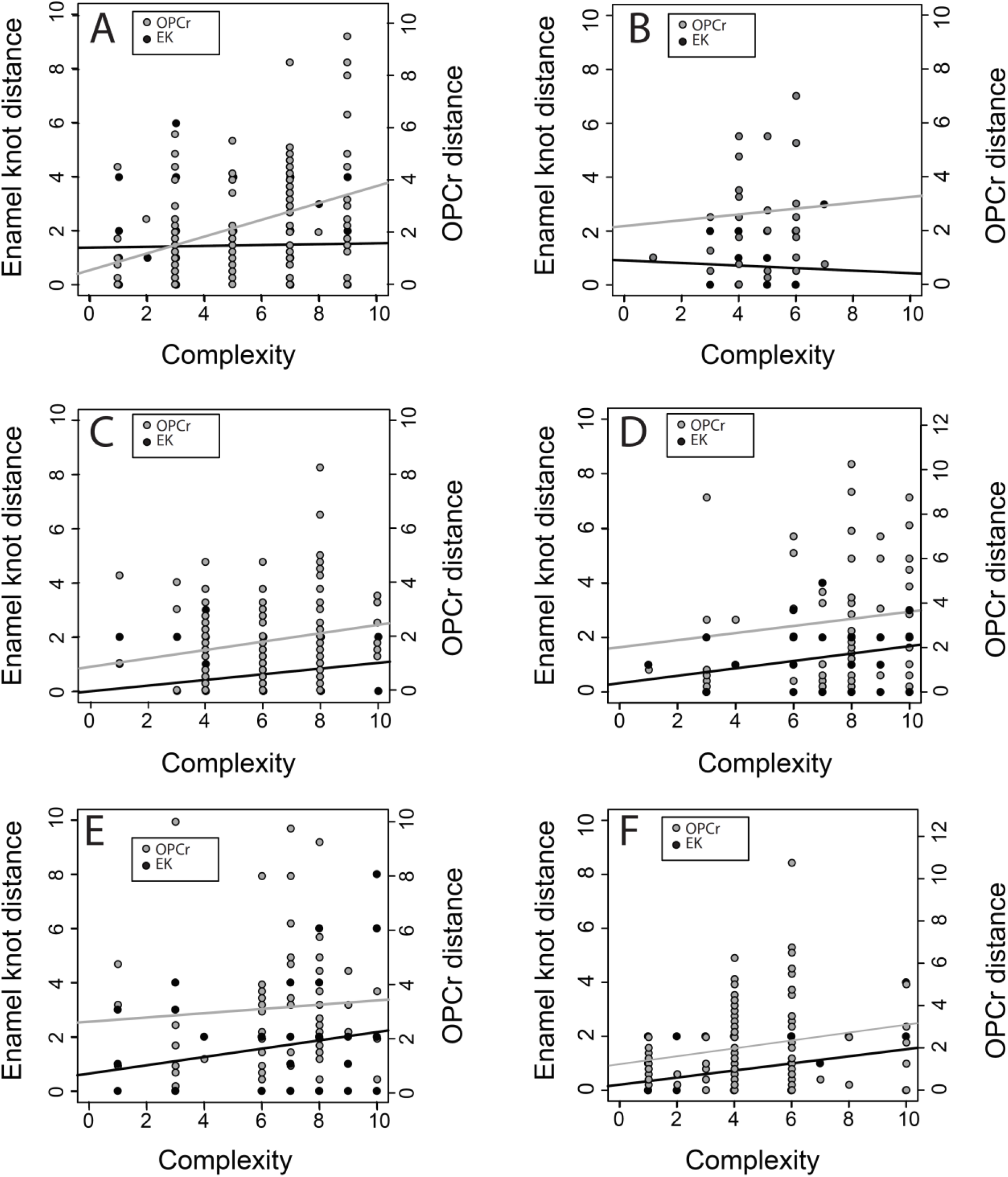
Relationship between tooth complexity and nearest-neighbour distance measured by enamel knot number or OPCr patches. (*A*) Activator autoactivation (Act) and activator diffusion rate (Da). (*B*) Activator autoactivation (Act) and inhibitor diffusion rate (Di). (*C*) Activator diffusion rate (Da) and inhibitor diffusion rate (Di). (*D*) Epithelial growth rate (Egr) and activator autoactivation (Act). (*E*) Epithelial growth rate (Egr) and activator diffusion rate (Da). (*F*) Epithelial growth rate (Egr) and inhibitor diffusion rate (Di).

**Table S1.** Pairwise parameter combinations used in ToothMaker simulations. Dimensionless parameter ranges and step size (brackets) are indicated.

| Developmental parameter | Inhibitor diffusion rate (Di) | Activator autoactivation (Act) | Activator diffusion rate (Da) |
| --- | --- | --- | --- |
| **Epithelial growth rate (Egr)** | Egr: 0.01–0.02 ($1.25 \times 10^{-3}$)<br>Di: 1–20 (1) | Egr: 0.01–0.02 ($1.25 \times 10^{-3}$)<br>Act: 0.1–0.125 ($5 \times 10^{-3}$) | Egr: 0.01–0.02 ($1.25 \times 10^{-3}$)<br>Da: 0.01–0.05 ($2.5 \times 10^{-3}$) |
| **Inhibitor diffusion rate (Di)** | – | Act: 0.1–0.125 ($2.5 \times 10^{-3}$)<br>Di: 1–15 (1) | Da: 0.01–0.05 ($5 \times 10^{-3}$)<br>Di: 1–15 (1) |
| **Activator autoactivation (Act)** | – | – | Act: 0.1–0.125 ($2.5 \times 10^{-3}$)<br>Da: 0.01–0.05 ($2.5 \times 10^{-3}$) |

